# A global pause generates nonselective response inhibition during selective stopping

**DOI:** 10.1101/2023.03.02.530898

**Authors:** Corey G. Wadsley, John Cirillo, Arne Nieuwenhuys, Winston D. Byblow

**Author notes:** Corresponding author:* Professor Winston Byblow.

## Abstract

1

Response inhibition is essential for terminating inappropriate actions. Selective response inhibition may be required when stopping part of a multicomponent action. However, a persistent response delay (*stopping-interference effect*) indicates nonselective response inhibition during selective stopping. This study aimed to elucidate whether nonselective response inhibition is the consequence of a global pause process during attentional capture or specific to a nonselective cancel process during selective stopping. We hypothesised that the stopping-interference effect would be larger in response to stop than ignore signals, owing to stronger nonselective response inhibition for explicit selective stopping. Twenty healthy human participants of either sex performed a bimanual anticipatory response inhibition paradigm with selective stop and ignore signals. Frontocentral and sensorimotor beta (β)-bursts were recorded with electroencephalography. Corticomotor excitability (CME) and short-interval intracortical inhibition (SICI) in primary motor cortex were recorded with transcranial magnetic stimulation. Behaviourally, responses in the non-signalled hand were delayed during selective ignore and stop trials. The response delay was largest during selective stop trials and indicated that the stopping-interference effect could not be attributed entirely to attentional capture. A stimulus-nonselective increase in frontocentral β-bursts occurred during stop and ignore trials, whilst sensorimotor response inhibition was reflected in maintenance of β-bursts and SICI relative to disinhibition observed during go trials. Signatures of response inhibition in the sensorimotor cortex contralateral to the responding hand were not associated with the magnitude of stopping-interference. Therefore, nonselective response inhibition during selective stopping results primarily from a nonselective pause process but does not entirely account for the stopping-interference effect.

**Significance statement:** Selective stopping is a complex form of response inhibition where a person must execute and cancel part of an action at the same time. A stopping-interference effect exemplifies the complexity of selective stopping. The present study examined whether nonselective response inhibition during selective stopping results from a global pause during attentional capture or is specific to a deliberate cancel process. Behaviourally, the interference effect was larger during selective stop stimuli than selective ignore stimuli. However, neurophysiological signatures of nonselective response inhibition were elicited in response to both stop and ignore stimuli. These findings indicate that nonselective response inhibition during selective stopping results primarily from a nonselective pause process but does not entirely account for the stopping-interference effect.

## 3 Introduction

Humans rely on response inhibition to navigate dynamic environments. Selective stopping can be required when complex behaviours are guided by various stimuli. Stimulus-selective stopping occurs when response inhibition is required for only a subset of salient stimuli (Bissett & Logan, 2014), for example, needing to stop for a red but not green traffic signal when driving. Response-selective stopping occurs when response inhibition is required for only part of a multicomponent action (Coxon et al., 2007), for example, needing to stop one hand from pressing keys while the other hand continues playing the piano. Enacting a selective response inhibition process to support stopping would provide optimal behaviour in these contexts. However, selective stopping is often supported by nonselective response inhibition, whereby stopping is momentarily engaged across all effectors in response to any salient stimulus (for a review, see Duque et al., 2017).

A persistent response delay (termed “stopping-interference effect”) occurs in non-stopped effectors during response-selective stopping (Wadsley et al., 2022c). The stopping-interference effect has been replicated across various response inhibition paradigms and indicates an underlying constraint on selective stopping (Aron & Verbruggen, 2008; Coxon et al., 2007). A restart model predicts that stopping-interference is caused by nonselective response inhibition and then a selective restart of non-stopped effectors (De Jong et al., 1995; MacDonald et al., 2017). Stimulus-selective stopping can also be nonselective, whereby response inhibition is engaged to stimuli indiscriminately while deciding to respond (Wessel & Aron, 2017). Stopping-interference can be influenced by nonselective response inhibition generated at the stimulus and response levels.

Response inhibition manifests in the sensorimotor cortex. Typical sensorimotor signatures of response inhibition are beta (β) band oscillations measured with electroencephalography (EEG) and corticomotor excitability (CME) measures from motor evoked potentials derived from transcranial magnetic stimulation (TMS) of primary motor cortex. Recent methodological advances have provided for β-burst analyses (Shin et al., 2017) that indicate unimanual response preparation and inhibition are marked by a decrease and then reinstatement of β-bursts, respectively (Wessel, 2020). Response inhibition is also marked by CME suppression (Jana et al., 2020), which may be driven by increased gamma-aminobutyric acid (GABA)_A_ receptor-mediated inhibition measured with paired-pulse TMS protocols of short-interval intracortical inhibition (SICI) (Coxon et al., 2006; Hermans et al., 2019). The above signatures of response inhibition have also been observed in the sensorimotor cortex contralateral to the responding hand (MacDonald et al., 2014; Wadsley et al., 2022b) and demonstrate a predominance of nonselective response inhibition during selective stopping (Raud et al., 2020). However, converging evidence has indicated that global motor suppression can occur during attentional capture of non-stopping stimuli (Iacullo et al., 2020; Wessel & Aron, 2017). Nonselective response inhibition during selective stopping may, in-part, be generated by global motor suppression during attentional capture.

Response inhibition during selective stopping may best be understood with a pause-then-cancel model. The model posits that stopping is achieved by a global pause process generated during attentional orienting to stimuli and a cancel process triggered on recognition of a stop-signal (Diesburg & Wessel, 2021; Shin et al., 2017). The specificity of response inhibition to the pause and cancel processes can be discerned by stimulus-selective stopping paradigms that include “ignore” signals and stop signals (e.g., Tatz et al., 2021). Ignore signals are presented similarly to stop signals but require the participant to continue the planned response rather than stop it (Sharp et al., 2010). Inhibition is likely a product of the pause process if observed during both stop and ignore signals, whereas inhibition exclusive to stop signals likely reflects the cancel process (Diesburg & Wessel, 2021). However, to date, no studies have experimentally examined to what extent pause and cancel processes contribute to nonselective response inhibition in a response-selective stopping context.

The present study aimed to elucidate the specificity of inhibitory signatures to pause and cancel processes during response-selective stopping (hereby referred to as “selective stopping”). Selective stopping was assessed using an anticipatory response inhibition (ARI) paradigm with partial-ignore and partial-stop trials. Frontocentral and sensorimotor signatures of response inhibition were assessed with EEG and TMS. Four primary hypotheses were tested: 1) Responses will be delayed more in the non-signalled hand during partial-stop than partial-ignore trials relative to a standard go context. 2) Post-signal frontocentral and sensorimotor β-bursts will increase nonselectively during partial-ignore and partial-stop trials. 3) Partial-ignore and partial-stop trials will be marked by nonselective suppression of corticomotor excitability after signal onset. 4) Post-signal short-interval intracortical inhibition (SICI) will be greater during partial-stop than partial-ignore and go trials in the signalled hand.

## 4 Methods

### 4.1 Participants

Twenty healthy adults volunteered to participate (11 female, 9 male; mean age 27.7 yrs., range 22 to 40 yrs.) All participants were right-handed (mean laterality quotient 0.81; range 0.5 to 1) as determined with the abbreviated Edinburgh Handedness Inventory (Veale, 2014). The study was approved by the University of Auckland Human Participants Ethics Committee (Ref. UAHPEC22709).

### 4.2 Task protocol

A multicomponent anticipatory response inhibition paradigm was used to assess selective stopping (Wadsley et al., 2022c). Participants were seated comfortably in front of an LG-24GL600F-B computer monitor (144 Hz refresh rate, ∼60 cm viewing distance) with their index fingers rested on blocks shoulder-width apart. Block height was adjusted to minimise postural activity observed from electromyography of the task-relevant *first dorsal interosseous* (FDI) muscle. A mechanical switch was positioned ∼1 cm above each block, so responses required sagittal index finger abduction. The task was designed with the Selective Stopping Toolbox (Wadsley et al., 2022a) integrated with PsychoPy software (v2022.1.4; Peirce et al., 2019) for accurate timing. A custom Arduino Leonardo response board was used to synchronise task equipment.

The default task display consisted of a grey background and two white bars (10 cm high, 1.5 cm wide). A black horizontal target line was positioned behind each bar at 80% of its total height. Each trial consisted of a fixation (0.5 – 1 s), trial (1.5 s), and feedback (1.2 s) period. Trial onset was signalled when the bars appeared to ‘fill’ from bottom to top. Each bar took 1 second to fill completely. Most trials were Go trials, where the objective was to cease the left and right bars from filling as close as possible to the respective target lines (target response time = 800 ms) by pressing the corresponding switches. In some trials, the left or right bar would change colour from black to cyan or magenta while filling as a means of providing partial ‘ignore’ and ‘stop’ signals (Figure 1A). Participants were instructed to execute the planned Go response on Partial-ignore trials or cancel the planned response on Partial-stop trials. The colour assignment was counterbalanced across participants and partial trials were equally distributed across the left and right bar presentations.

**Figure 1.**
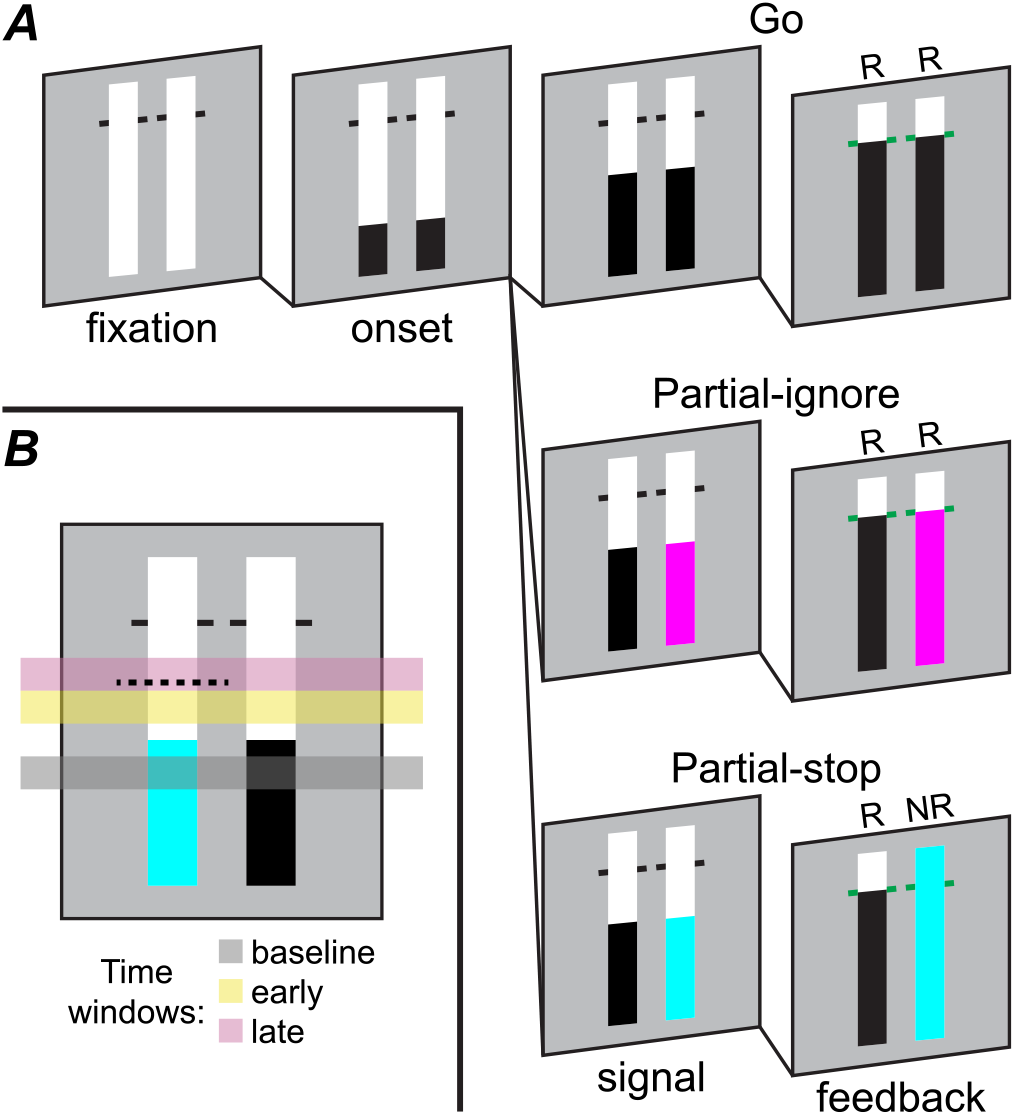
A stimulus-selective anticipatory response inhibition task. **A**: Trials began with two empty bars. Trial onset was signalled by both bars starting to fill from bottom to top synchronously. Most trials were go trials, where the objective was to respond (R) with both hands as close as possible to the target response time (800 ms). During partial trials, either the left or right bar could change colour prior to the target response time (*right bar presentation depicted*). Participants were instructed to either ignore the colour change and respond as normal (partial-ignore trial) or withhold the response (NR) in the signalled hand (partial-stop trial) based on the colour change. **B**: A representative partial-stop_left_ trial indicating the time windows of interest relative to partial-signal onset (i.e., colour change) for the EEG task component. Horizontal dashed line indicates the post-signal stimulus time for the left hand during the TMS task component.

**Figure 2.**
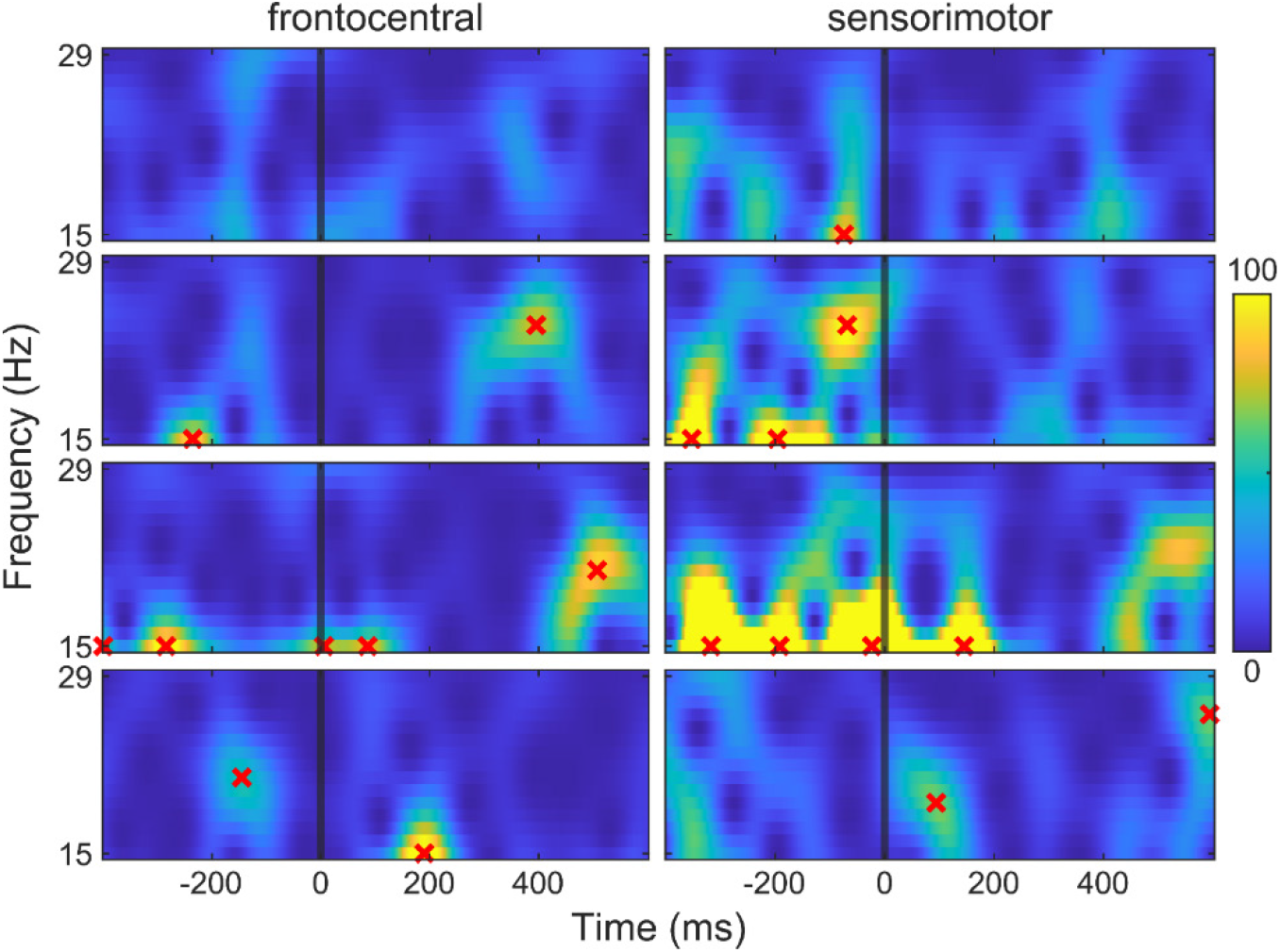
β-band epochs from frontocentral (electrode FCz) and sensorimotor (electrodes C3/C4) regions from four randomly selected trials in a representative participant. Epochs are locked to partial-signal onset (-400 to 600 ms). Red crosses indicate the automatically detected β-bursts. Calibration bar represents power in mV^2^.

**Figure 3.**
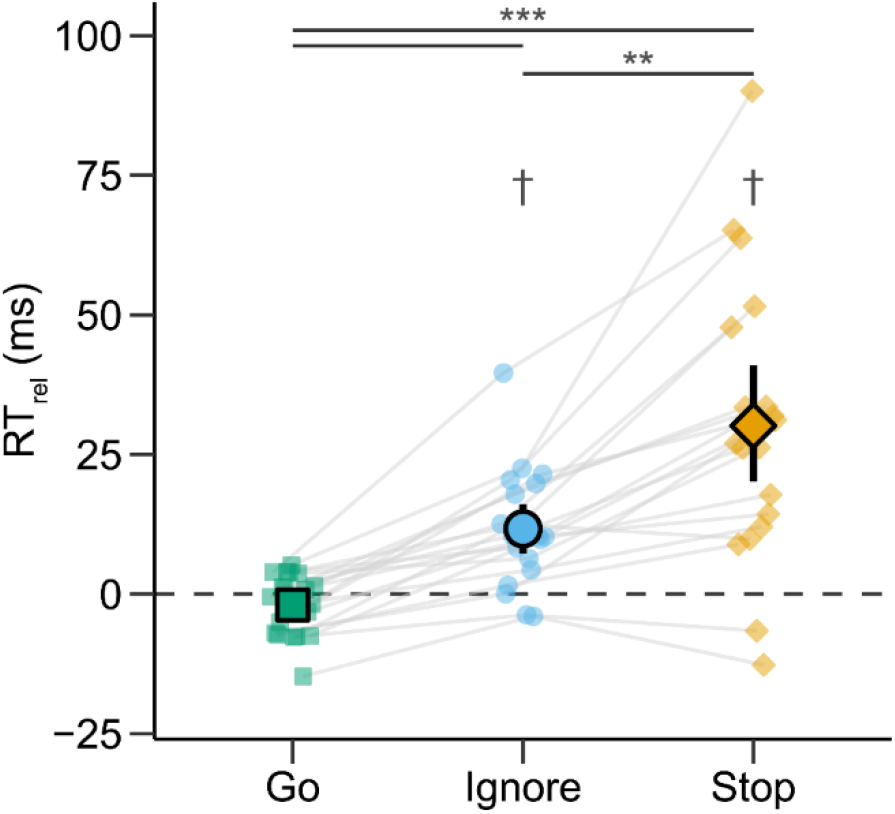
Relative response times (RT_rel_) during go, partial-ignore, and partial-stop trials. RT_rel_ is calculated by subtracting the target response time (800 ms, *dashed line*) from mean RT, where values greater than 0 indicate late responses relative to the target. RT_rel_ was calculated from the non-signalled hand for partial trials. Point ranges represent means with 95% bootstrap confidence intervals. Posterior odds: ** > 10, *** > 100; † > 100 for one-sample t-test against 0.

**Figure 4.**
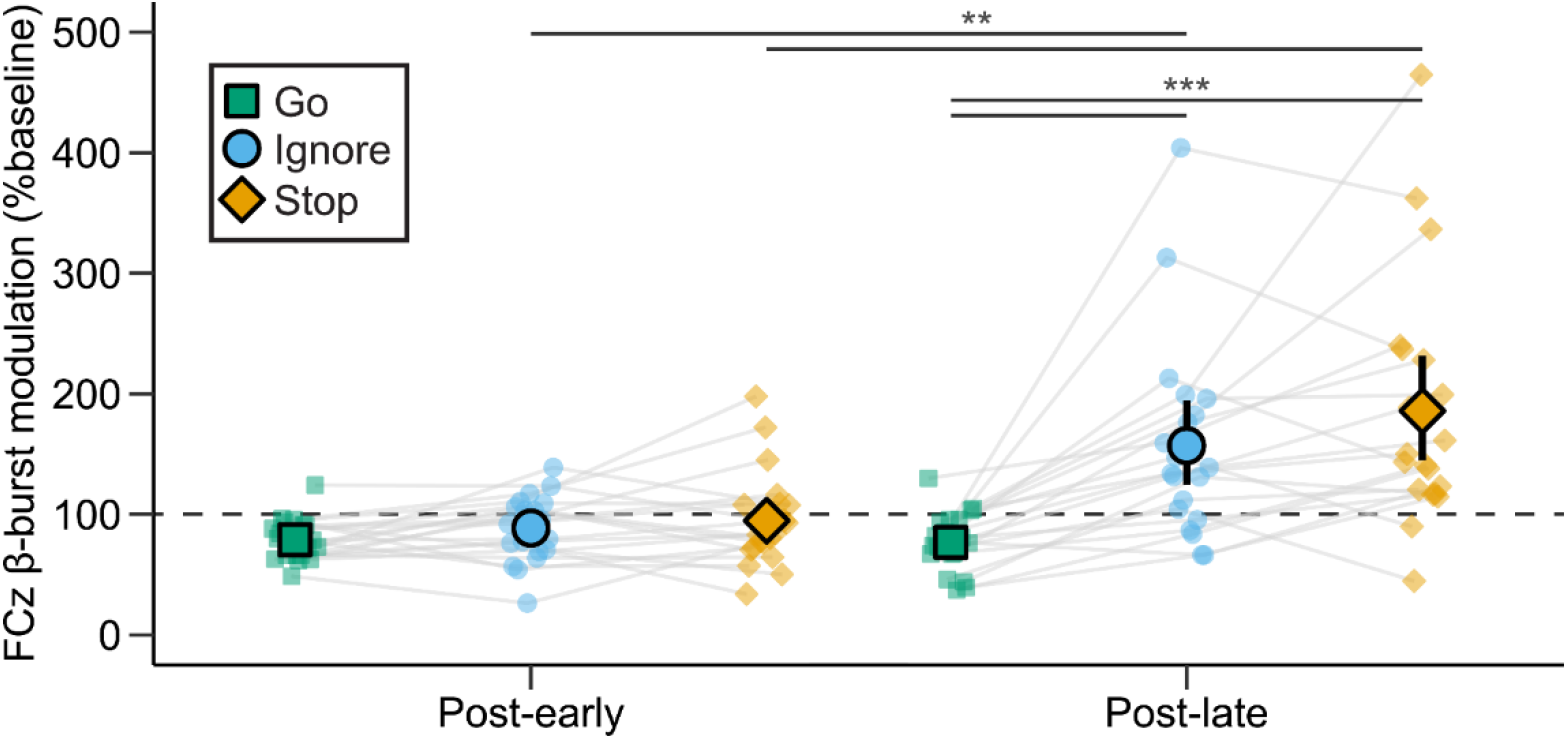
Frontocentral (electrode FCz) β-burst modulation at early (50 to 150 ms) and late (150 to 250 ms) post-signal time windows during go, partial-ignore, and partial-stop trials. Modulation was calculated as the percent change in β-bursts from a pre-signal baseline (-150 to -50 ms, *dashed line*) time window, where values greater than 100 indicate more β-bursts. Point ranges represent means with 95% bootstrap confidence intervals. Posterior odds: ** > 10, *** > 100.

**Figure 5.**
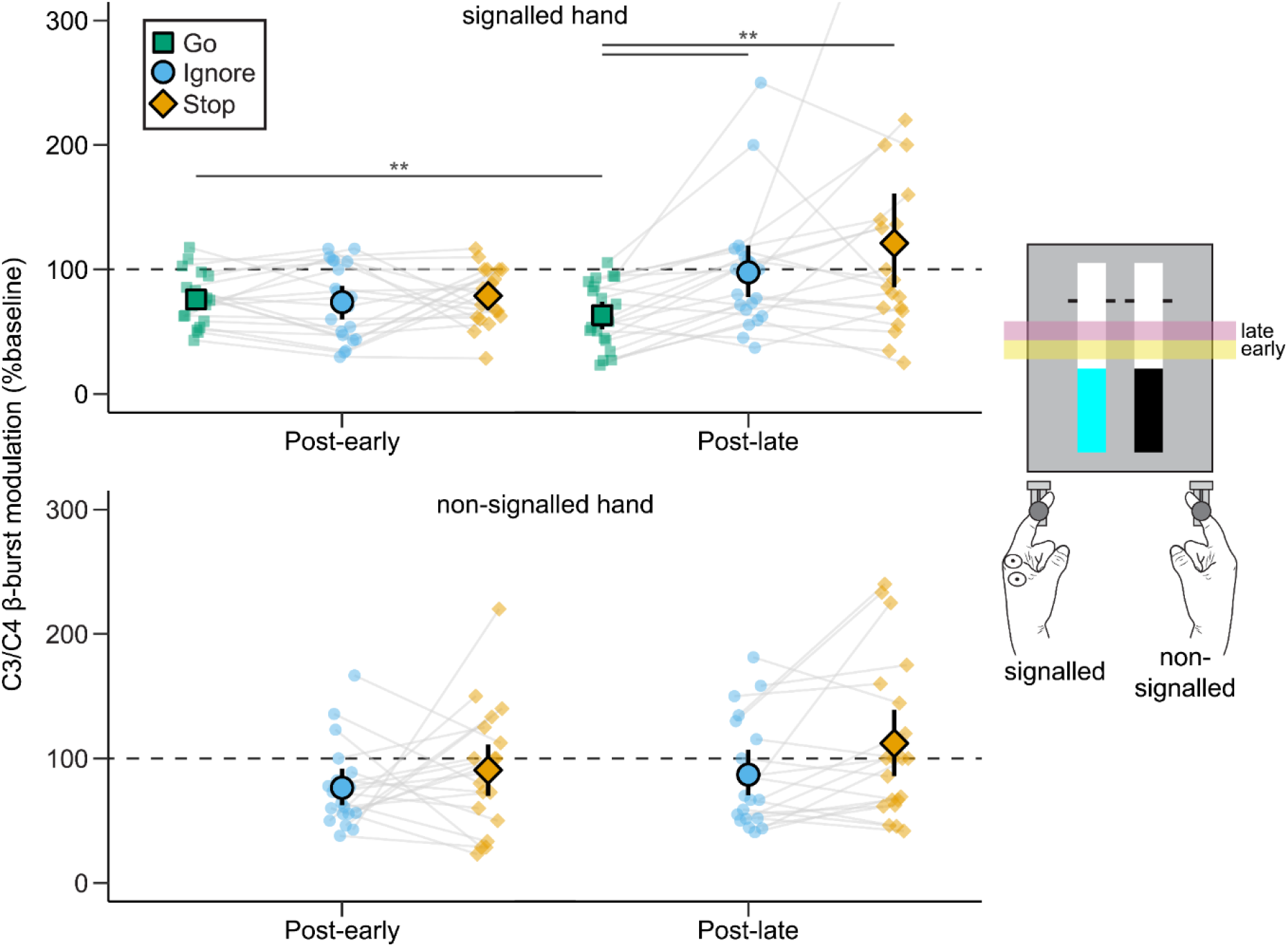
Sensorimotor (electrodes C3/C4) β-burst modulation at early (50 to 150 ms) and late (150 to 250 ms) post-signal time windows during go, partial-ignore, and partial-stop trials. β-burst modulation was calculated for the sensorimotor cortex contralateral to the signalled (*top*) and non-signalled hand (*bottom*), as indicated by an example partial-stop_left_ trial. Modulation is calculated as the percent change in β-bursts from a pre-signal baseline (-150 to -50 ms, *dashed line*) time window, where values greater than 100 indicate more β-bursts. Point ranges represent means with 95% bootstrap confidence intervals. Posterior odds: ** > 10.

**Figure 6.**
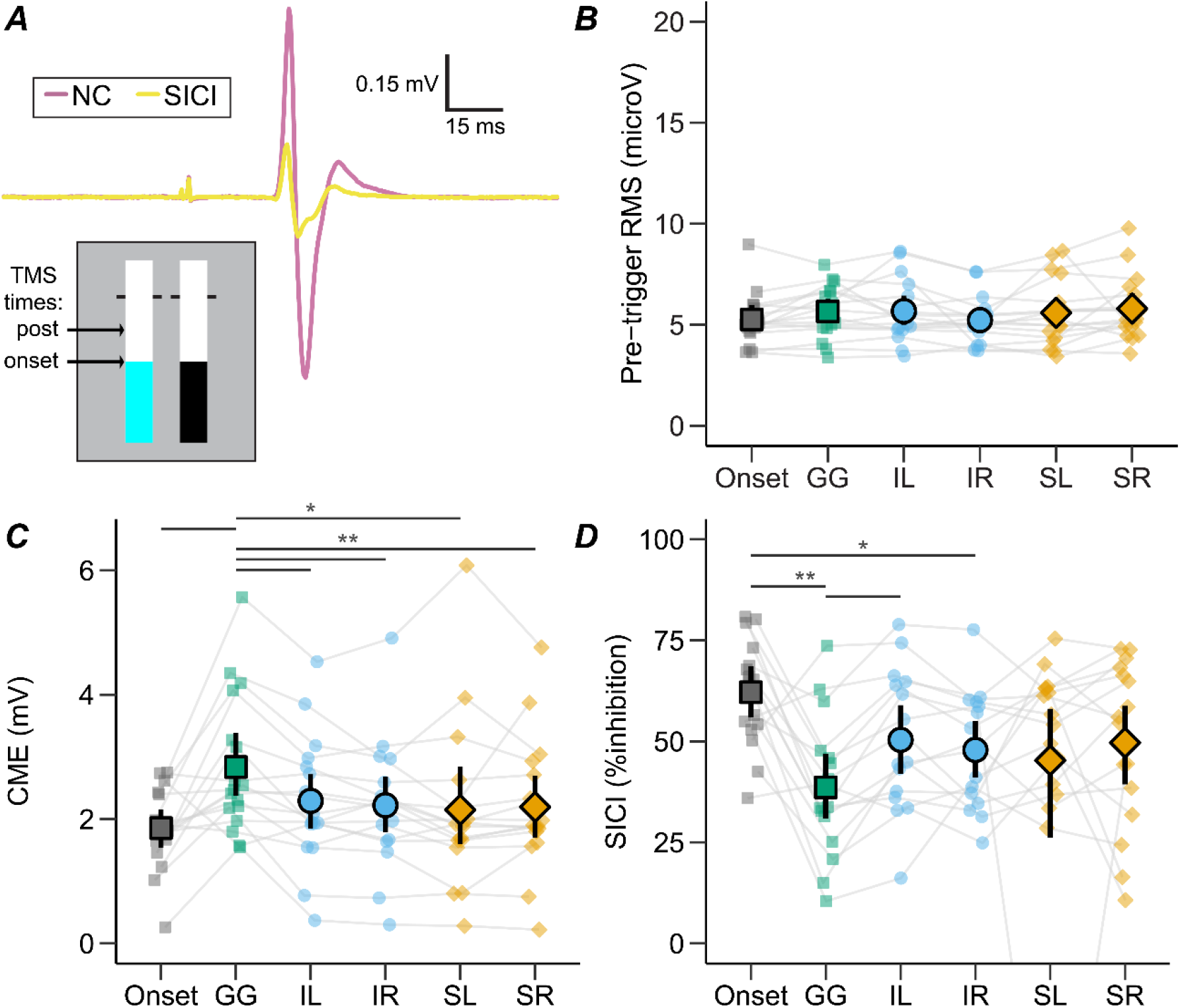
Transcranial magnetic stimulation (TMS) measures from the left hand at onset or post-signal in go (GG), partial-ignore_left_ (IL), partial-ignore_right_ (IR), partial-stop_left_ (SL), and partial-stop_right_ (SR) trials. TMS was delivered at or 175 ms after the participant-specific stop-signal delay for onset or post-signal stimuli, respectively. **A**: Electromyography traces of motor evoked potentials (MEPs) during nonconditioned (NC) and short-interval intracortical inhibition (SICI) stimulation protocols at rest. Each trace is the average of 25 trials recorded from the task-relevant first dorsal interosseous muscle in a representative participant. For SICI, a subthreshold conditioning stimulus was delivered 2 ms prior to the test stimulus. **B**: Pre-trigger root mean square (RMS) muscle activity across stimulation types. **C**: Corticomotor excitability (CME) calculated as mean NC peak-to-peak MEP amplitude. **D**: SICI calculated as percent inhibition, where greater values indicate smaller conditioned relative to NC MEP amplitude (i.e., more inhibition). Point ranges represent means with 95% bootstrap confidence intervals. Posterior odds: * > 3, ** > 10.

The stop-signal delay (SSD) was initially set so that the colour change occurred 250 ms prior to the target and was then adjusted in steps of 35 ms (∼5 frames) based on partial-stop trial success. The SSD was increased (i.e., less time before the target) after successful stopping and decreased (i.e., more time before the target) after unsuccessful stopping to obtain an average stop success of ∼50%. The SSD during partial-ignore trials was based on the most recent value from partial-stop trials to ensure equivalent presentation times. Points were awarded based on RT and stop success for the left and right response to encourage high accuracy. The number of points earned was signalled by changing the colour of the target lines during the intertrial interval (green [100 points]: < 25 ms or successful stop; yellow [50 points]: 26 – 50 ms: orange [25 points]: 51 – 75 ms; red [0 points]: > 75 ms or failed stop).

The experiment was split into EEG and TMS components. Task instructions were given at the experiment start, after which participants completed a practice block of go and partial-ignore trials and then of go, partial-ignore, and partial-stop trials for familiarisation. The EEG component started after familiarisation and was followed by the TMS component after a ∼15-minute break. The task for each component consisted of 600 trials (400 go, 50 partial-ignore_left_, 50 partial-ignore_right_, 50 partial-stop_left_, 50 partial-stop_right_) evenly split across eight blocks of 75 trials with a random order. Participants were informed that their primary goal was to earn as many points as possible for each component. The current block and total score were updated and displayed to the participant between blocks. The entire experiment lasted 3 hours on average.

### 4.3 EEG data acquisition and preprocessing

Continuous scalp-EEG was recorded using 32-channel Acticap Ag/Cl active shielded electrodes and BrainProducts MRPlus amplifiers at a sampling rate of 2500 Hz and resolution of 0.1 µV. The ground and online reference electrodes were placed at AFz and Cz, respectively. Before data collection, a predetermined electrode impedance (≤20 kΩ) was achieved for each channel. Continuous EEG data were recorded in Brain Vision Recorder (BrainProducts GmbH) and stored for later offline analysis.

EEG data were preprocessed using EEGLAB in MATLAB (Delorme & Makeig, 2004), as performed in Wadsley et al. (2022b). Data were low-pass filtered using a hamming window filter with a 6-dB cut-off at 45 Hz (10 Hz transition bandwidth), resampled to 512 Hz, and then high-pass filtered using a hamming window filter with a 6 dB cut-off at 0.5 Hz (1 Hz transition bandwidth). Filtered data were then epoched around trial onset (-1 to 2 s) and visually inspected for channel-wise and then epoch-wise artefacts. Channels (*M* = 0.4 out of 32 channels) and epochs (*M* = 3.9%) with excessive noise or irregular artefacts were manually rejected. An independent component analysis was then performed on the remaining data to remove components (*M* = 3.1 components) reflecting regular artefacts (e.g., eye movements, muscle activity) semiautomatically using ICLabel (Pion-Tonachini et al., 2019). Previously removed channels were then replaced using spherical spline interpolation. Lastly, data were transformed into a reference-free montage using current source density interpolation to reduce the effect of volume conduction (Tenke & Kayser, 2005).

β-burst detection was adapted from Wessel (2020) and quantified for frontocentral (electrode FCz) and sensorimotor (electrodes C3 & C4) regions of interest. Preprocessed EEG data were first epoched relative to partial signal onset (-1 to 1.2 s). The average SSD during partial trials was used to epoch go trials (Jana et al., 2020). Signal-locked epochs were then convolved with a complex Morlet wavelet for 15 evenly spaced frequencies within the β-band (15 – 29 Hz). Time-frequency power estimates were calculated by taking the squared magnitude of the convolved data. Local maxima within the time-frequency epochs were then determined using the MATLAB function *imregionalmax* from -400 to 600 ms relative to signal onset. A β-burst was defined as local maxima within an epoch that exceeded 6× the median for the electrode of interest (Enz et al., 2021; Wessel, 2020) (Figure 8.2).

### 4.4 Transcranial magnetic stimulation

Single- and paired-pulse TMS were delivered using a 70 mm figure-of-eight coil (100 µs pulse width) connected to a Magstim BiStim^2^ stimulator (The Magstim Company Ltd, Whitland). The coil was oriented to induce a posterior-to-anterior flowing current direction in the underlying cortical tissue. The optimal coil position for eliciting a motor evoked potential (MEP) in the left *first dorsal interosseous* (FDI) muscle was assessed and marked on the scalp. MEPs were recorded with surface electromyography (EMG). Ag-AgCl surface electrodes (CONMED) were positioned over left FDI in a belly-tendon montage, and a ground electrode was positioned on the posterior surface of the left hand. EMG activity was amplified (×1000), bandpass filtered (10 – 1000 Hz), and sampled at 2500 Hz with a CED interface system (MICRO1401mkII; Cambridge Electronic Design). EMG data were recorded in Signal software (v7.05) and stored for later offline analysis in MATLAB.

Motor thresholds for left FDI were determined using a maximum-likelihood parameter estimation by sequential testing strategy (Awiszus, 2011). Active motor threshold (AMT) was defined as the minimum stimulator output required to elicit an MEP amplitude of at least 0.2 mV during low-intensity voluntary contraction. Test stimulus (TS) intensity was determined as the minimum stimulator output required to elicit an MEP amplitude of at least 1 mV at rest. A subthreshold conditioning stimulus (CS) was delivered 2 ms prior to the test stimulus to assess SICI (Peurala et al., 2008). The CS intensity was first set to 70% AMT and, if necessary, adjusted to produce approximately 50% inhibition at rest. For all participants, CS intensity was below 100% AMT and remained constant throughout the task protocol.

TMS was delivered at rest and during the task. Nonconditioned and SICI measures at rest were collected over 2 blocks of 25 trials with a random stimulation order. Stimulated trials were intermixed with non-stimulated trials during the TMS task component. SSD for Partial-ignore and Partial-stop trials during the TMS task was fixed and set to the participant-specific stopping time (Hermans et al., 2019): *SSD*_*TMS*_ = 800 *ms* − [*GoRT*_*EEG*_ − *SSD*_*EEG*_] − 50 *ms*, where GoRT_EEG_ and SSD_EEG_ reflect the mean values from the EEG task. In-task TMS was delivered at SSD_TMS_ or 175 ms after SSD_TMS_ to investigate the influence of post-signal inhibitory signatures. The late stimulation time point was selected to coincide with the nonselective dip observed in CME of the non-signalled hand during similar response inhibition tasks (Cowie et al., 2016; MacDonald et al., 2014). A total of 25 nonconditioned (TS only) and 25 conditioned (CS and TS) stimulations were collected at SSD onset and the late stimulation timing for go, partial-ignore, and partial-stop trials. In the case of partial trials, TMS measures were collected from the left hand when the partial signal was presented on the left (signalled) and right (non-signalled) sides separately. In total, there were 300 stimulated and 300 non-stimulated trials during the TMS task.

### 4.5 Dependent measures

#### 4.5.1 Behavioural data

Behavioural data were analysed with custom scripts in Python and taken only from the EEG session to avoid the influence of TMS on performance. Success was determined as the percentage of trials with correct responses within the trial period for go (both hands respond), partial-ignore (response in signalled hand) and partial-stop (no-response in signalled hand) trials. Relative RT (RT_rel_) was calculated by subtracting the target RT from the mean RT across the hands for each trial type. In the case of partial-ignore and partial-stop trials, RT was taken from the hand to which the partial signal was not presented. Negative and positive RT_rel_ values indicate early and late responses relative to the target. Mean SSD for partial trials was made relative to the target, where values closer to 0 indicate less time for stopping.

#### 4.5.2 EEG data

β-bursts were determined for frontocentral and sensorimotor regions of interest. The number of β-bursts was determined for each trial type within three time periods: a baseline period before the partial signal (-150 to 50 ms), as well as early (50 to 150 ms) and late (150 to 250 ms) periods after the partial signal (see Figure 8.1B). β-burst proportion was calculated as the sum of β-bursts divided by the number of trials for each trial type and time separately. For sensorimotor β-bursts, the proportion was calculated separately for the sensorimotor cortex contralateral to the signalled and non-signalled hand during partial trials. β-burst modulation was then calculated as the percent change in proportion from baseline to the early and late time windows, where values greater than 100 indicate an upregulation of β-bursts.

#### 4.5.3 TMS data

Peak-to-peak MEP amplitude was calculated for the left FDI between 20 and 40 ms post-stimulation. MEPs were excluded when pre-trigger root mean square EMG activity exceeded 20 µV in a -55 to -5 ms pre-trigger window (*M* = 3.6% of stimulated trials). The top and bottom 10% of MEP amplitudes were trimmed for each conditioned before calculating mean MEP amplitude (Wilcox, 2010). Corticomotor excitability (CME) was determined as the mean nonconditioned (NC) MEP amplitude for each condition. The magnitude of SICI was calculated for each condition as: 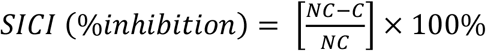, where C and NC refer to each participant’s mean conditioned and nonconditioned MEP amplitude respectively. In this case, values greater than 0 indicate an inhibitory effect of the SICI protocol.

### 4.6 Statistical analyses

Data were analysed with Bayesian analyses of variance (ANOVA) using JASP software (Version 0.16.3; JASP Team, 2020). All models included random slopes and were fitted across 100,000 iterations with participant modelled as a random intercept (van den Bergh et al., 2022). Normality of data and model-averaged residual plots were visually inspected before ANOVA (van den Bergh et al., 2020). Logarithmic transformations were used for non-normal data. Interaction effects were determined by comparing models with the interaction term against matched models without the term (van den Bergh et al., 2020).

Evidence for main effects and interactions were determined using Bayes factor in favour of the alternative hypothesis (BF_10_ ± percent error), where values greater than 1 indicate support for the alternative hypothesis and values less than 1 support the null hypothesis. The strength of evidence was determined using a standard BF_10_ classification table (BF_10_ < 0.3: moderate evidence for null hypothesis; 0.3 ≤ BF_10_ ≤ 3: inconclusive evidence; BF_10_ > 3: moderate evidence for alternative hypothesis) (van Doorn et al., 2021). Post-hoc pairwise comparisons were performed using Bayesian paired t-tests when a main effect or interaction was found.

Corrected posterior odds (*O*_post._) were calculated by multiplying the uncorrected BF_10_ by the adjusted prior odds (van den Bergh et al., 2020). Prior odds were adjusted using the Westfall multiple comparisons approach (Westfall, 1997). Data are presented as non-transformed mean ± standard deviation unless otherwise specified.

RT_rel_ was assessed with a one-way ANOVA with the factor of Trial Type (go, partial-ignore, partial-stop) to test the first hypothesis that response delays will be larger during partial-stop trials than during partial-ignore and go trials. In this case, response times were taken from the non-signalled hand or averaged across the hands for partial and go trials, respectively. One-sample t-tests against 0 were used to test RT_rel_ and verify stopping-interference during partial trials. A Bayesian paired t-test was also used to verify that SSD was not meaningfully different between partial-ignore and partial-stop trials.

Frontocentral and sensorimotor β-burst modulation was assessed with two-way ANOVAs with the factors of Trial Type (go, partial-ignore, partial-stop) and Time (early, late) to test the second hypothesis that post-signal β-bursts will increase nonselectively during partial trials. To test the selectivity of response inhibition, sensorimotor β-burst modulation for the non-signalled hand was assessed with a one-way ANOVA with the factor of Trial Type (go, partial-ignore, partial-stop). One-way ANOVAs with the factor of Trial Type were used to verify that β-burst proportion at frontocentral and sensorimotor regions did not differ between trial types at baseline. A two-way ANOVA with the factors of Region (frontocentral, sensorimotor) and Burst (with, without) was used to determine whether the presence of β-burst within the early or late period was associated with more stopping-interference.

Sensorimotor β-burst presence was taken from the cortex contralateral to the non-signalled hand. To test the third and fourth hypotheses, that partial trials will be marked by nonselective suppression of CME after partial signal onset and that post-signal SICI will be greatest during partial-stop trials in the signalled hand, CME and SICI were analysed with a one-way ANOVA with the factor of Stimulation (onset, post-go, post-ignore_left_, post-ignore_right_, post-stop_left_, post-stop_right_). Pre-trigger root-mean square (RMS) activity was also assessed with a one-way ANOVA with the factor of Stimulation (onset, post-go, post-ignore_left_, post-ignore_right_, post-stop_left_, post-stop_right_) to verify that modulation in background EMG activity did account for CME or SICI results from the analyses above.

## 5 Results

### 5.1 Behaviour

Participants performed the task correctly, as indicated by high average success for go (99.3 ± 1.0 %) and partial-ignore (87.2 ± 9.8 %) trials and success close to 50% for partial-stop trials (47.9 ± 1.9 %). For RT_rel_ (Figure 8.3), there was a main effect of Trial Type (1.58 × 10_7_ ± 0.41%). RT_rel_ during partial-stop trials (30.13 ± 24.68 ms) was greater than go (−2.00 ± 5.39 ms; *O*_post._ = 3.04 × 10_3_) and partial-ignore trials (11.68 ± 10.26 ms; *O*_post._ = 87.51), which also differed from each other (*O*_post._ = 1.47 × 10_4_). One-sample t-tests against 0 supported the presence of an interference effect in partial-ignore (*O*_post._ = 478.22) and partial-stop trials (*O*_post._ = 996.31) but not go trials (*O*_post._ = 0.10). There was no meaningful evidence for a difference between SSD during partial-ignore (−306.70 ± 48.55 ms) and partial-stop trials (−308.55 ± 49.70 ms; *O*_post._ = 0.50).

### 5.2 Frontocentral and sensorimotor β-bursts

Baseline frontocentral β-burst proportion (0.19 ± 0.07 baseline bursts per trial) was not meaningfully different between go, partial-ignore and partial-stop trials (BF_10_ = 0.35). For post-signal frontocentral β-burst modulation (Figure 8.4), there was a Trial Type × Time interaction (BF_10_ = 3.76 × 10^4^ ± 4.3%). There was an increase in frontocentral β-burst modulation from early to late for partial-ignore (early = 88.6 ± 27.2 %, late = 157.0 ± 82.9 %; *O*_post._ = 81.43) and partial-stop trials (early = 94.7 ± 40.0 %, late = 185.8 ± 102.4 %); *O*_post._ = 77.57), but inconclusive evidence for go trials (early = 78.8 ± 17.2 %, late = 76.7 ± 23.8 %; *O*_post._ = 0.36). There were no differences between trial types at the early time point (all *O*_post._ ≤ 0.35), whereas frontocentral β-burst modulation at the late time point was greater than go trials for partial-ignore (*O*_post._ = 5.06 × 10^3^) and partial-stop trials (*O*_post._ = 1.04 × 10^4^). There was inconclusive evidence for a difference between partial-ignore and partial-stop trials (*O*_post._ = 0.38).

Baseline sensorimotor β-burst proportion (0.25 ± 0.12 baseline bursts per trial) was not meaningfully different between go, partial-ignore and partial-stop trials (BF_10_ = 0.50). For β-burst modulation in the sensorimotor cortex contralateral to the signalled hand (Figure 8.5), there was a Trial Type × Time interaction (BF_10_ = 298.16 ± 0.97%). There was a decrease in sensorimotor β-burst modulation from early to late for go (early = 75.9 ± 20.7 %, late = 63.3 ± 26.0 %; *O*_post._ = 38.06), but inconclusive evidence for partial-ignore (early = 73.7 ± 30.8 %, late = 97.6 ± 50.8 %; *O*_post._ = 2.18) and partial-stop trials (early = 78.8 ± 22.5 %, late = 121.1 ± 91.0 %; *O*_post._ = 0.86).

There were no differences between trial types at the early time point (all *O*_post._ ≤ 0.26). Sensorimotor β-burst modulation at the late time point was less during go trials than during partial-ignore (*O*_post._ = 66.85) and partial-stop trials (*O*_post._ = 29.84). There was inconclusive evidence for a difference between partial-ignore and partial-stop trials (*O*_post._ = 0.31). For β-burst modulation in the sensorimotor cortex contralateral to the non-signalled hand (Figure 8.5), there was inconclusive evidence for a main effect of Trial Type (BF_10_ = 0.40 ± 1.91%) or Time (BF_10_ = 0.71 ± 1.60%), nor for a Trial Type × Time interaction (BF_10_ = 0.42 ± 2.36%).

For stopping-interference during trials with (31.0 ± 22.6 ms) and without (32.8 ± 24.3 ms) β-bursts, there was a null main effect of Region (BF_10_ = 0.29 ± 1.55%) and inconclusive evidence for a main effect of Burst (BF_10_ = 0.41 ± 1.14%) and a Region × Burst interaction (BF_10_ = 0.44 ± 2.24%).

### 5.3 CME and SICI results

Three participants were excluded from the TMS component due to high TS intensities, which caused coil overheating during task blocks. After exclusions, data from 17 participants were available for TMS analyses. For the remaining participants, TS intensity was 60.0 ± 14.0% of maximum stimulator output, and CS intensity was equivalent to 83.2 ± 6.3% of active motor threshold. Participant-specific SSD during the TMS task was, on average, -356.5 ± 47.5 ms. On average 23.5 ± 1.1 trials were available for each TMS trial type before trimming. One-sample t-tests on the behavioural differences indicated that RT_rel_ was 2.6 ± 4.0 ms faster during non-stimulated trials in the TMS session than in the EEG session (*O*_post._ = 5.91). RT_rel_ differences between the sessions did not exceed 10 ms in any participant (range -8.6 to 6.2 ms), indicating that both sessions were comparable behaviourally. Figure 8.6A shows representative EMG traces from one participant. Pre-trigger RMS (Figure 8.6B) was below 10 µV for all participants (grand average: 5.53 ± 0.2 µV), with inconclusive evidence for a main effect of Stimulation (BF_10_ = 0.38 ± 0.11%).

For CME (Figure 8.6C), there was a main effect of Stimulation (BF_10_ = 2.35 × 10_3_ ± 0.13%). There was an increase in CME from onset (1.85 ± 0.65 mV) to post-go (2.84 ± 1.12 mV; *O*_post._ = 6.00) but not post-stop_left_ (2.15 ± 1.34 mV; *O*_post._ = 0.11) or post-stop_right_ (2.20 ± 1.07 mV; *O*_post._ = 0.23) trials, whilst evidence was inconclusive for post-ignore_left_ (2.29 ± 1.03 mV; *O*_post._ = 0.51) and post-ignore_right_ (2.22 ± 1.02 mV; *O*_post._ = 0.31). CME at the post go stimulation was greater than all partial trials (all *O*_post._ ≥ 7.52), which did not differ from each other (all *O*_post._ ≤ 0.3) except for inconclusive evidence between partial-ignore_left_ and partial-stop_right_ (*O*_post._ = 0.35).

One participant was excluded from SICI analyses because inhibition could not be elicited at rest reliably. For SICI (Figure 8.6D), there was a main effect of Stimulation (BF_10_ = 3.21 ± 0.08%). There was a decrease in SICI from onset (62.2 ± 13.3 %) to post-go (38.6 ± 17.2 %; *O*_post._ = 19.25) and post-ignore_right_ (47.9 ± 14.2 %; *O*_post._ = 3.67), but not post-stop_left_ (45.3 ± 35.5 %; *O*_post._ = 0.25), whilst evidence was inconclusive for post-ignore_left_ (50.4 ± 17.2 %; *O*_post._ = 0.85) and post-stop_right_ (49.7 ± 20.4 %; *O*_post._ = 1.01) trials. SICI at the post-go stimulation was less than the post-ignore_left_ stimulation (*O*_post._ = 32.2), while there were no differences between partial trials (all *O*_post._ ≤ 0.10).

## 6 Discussion

The present study provides novel insight into response inhibition during selective stopping. Overall, the findings indicate that common signatures of inhibitory control measured with EEG and TMS during selective stopping reflect a global pause rather than a selective cancel process. Below we discuss the main findings in relation to the primary hypotheses specifically and the selectivity of response inhibition more generally.

### 6.1 More interference during partial-stop than partial-ignore trials

In support of the first hypothesis, responses were more delayed during partial-stop than partial-ignore trials. Responses during go trials were close to the target and indicated that participants were not intentionally slowing to improve partial trial success (Verbruggen et al., 2013). Responses were delayed relative to go trials for partial-ignore and partial-stop trials, indicating that a nonselective (global) pause can generate interference during attentional capture. However, the interference effect was twice as large during partial-stop (∼32 ms) than partial-ignore (∼14 ms) trials, corroborating previous observations with the stop-signal task (Ko & Miller, 2013). Prolonged interference during partial-stop trials indicates that part of the constraint on selective stopping is specific to the explicit requirement to stop.

### 6.2 Signal-nonspecific increase in frontocentral β-bursts during partial trials

In support of the second hypothesis, frontocentral β-bursts were increased relative to baseline during partial-stop trials. Increased frontocentral β-bursts have been observed during unimanual stopping and are proposed to reflect signalling from upstream prefrontal regions to engage response inhibition (Hannah et al., 2020; Wessel, 2020). Frontocentral β-bursts increased only at the late period of interest (150 to 250 ms post signal), coinciding with the typical temporal progression of response inhibition (Jana et al., 2020). Interestingly, frontocentral β-bursting was also observed during partial-ignore trials, despite no explicit stopping requirement. Importantly, there was no modulation of frontocentral β-bursts during go trials where an additional signal was not presented. Overall, the stimulus-nonselective upregulation of frontocentral β-bursts provides evidence for a prevailing view that global motor suppression is engaged in response to infrequent stimuli (Iacullo et al., 2020). To the best of our knowledge, this is the first demonstration of a stimulus-nonselective increase in frontocentral β-bursts during selective stopping.

The modulation of sensorimotor β-bursts was also similar between partial-ignore and partial-stop trials. Sensorimotor β-bursts decreased from the early to the late period of interest during go trials, reflecting the expected disinhibition during response preparation (Kilavik et al., 2013; Wessel, 2020). In contrast, sensorimotor β-bursts were maintained during partial-ignore and partial-stop trials, which were both greater than go trials at the late period of interest. Functionally, maintenance of sensorimotor β-bursts may raise the response threshold to ensure that the planned go response is not enacted during stimulus processing (Muralidharan et al., 2022). A similar pattern of β-burst maintenance was observed in the sensorimotor cortex contralateral to the non-signalled hand, indicating that the pause process generated nonselective response inhibition. The stopping-interference effect was not larger during partial-stop trials with a frontocentral or a sensorimotor β-burst than those without a burst. Speculatively, the magnitude of the stopping-interference effect may depend more on how an individual prepares to go (Raud et al., 2020; Xu et al., 2015) rather than how selectively they stop. Overall, the stimulus-nonselective maintenance of sensorimotor and frontocentral β-bursts supports the predominance of a pause process during selective stopping.

### 6.3 Nonselective CME and SICI modulation indicate selective stopping

In support of the third hypothesis, nonselective CME suppression occurred during partial trials. As expected, a CME increase occurred during response preparation of go trials (Cowie et al., 2016; MacDonald et al., 2014) but not after signal onset during partial trials. CME was greater during go trials compared with partial-ignore and partial-stop trials, irrespective of whether the partial signal was presented to the stimulated left or non-stimulated right hand. Similar CME suppression of non-stopped effectors to ignore and stop signals has been observed during a unimanual stop-signal task (Tatz et al., 2021; Wessel & Aron, 2013). Interestingly, the stimulation time used in the current study (175 ms post signal) was the first point identified by Tatz et al. (2021), where CME differed between stop and ignore trials. A similar discrepancy between partial-ignore and partial-stop trials may emerge later during selective stopping due to a slower but more selective cancel process (Aron, 2011). While the above interpretation is speculative, the pattern of results indicates that the stimulus-nonselective pause process likely drives CME suppression early after a partial signal.

The fourth hypothesis was not supported since SICI modulation was not specific to partial-stop trials. In a unimanual stopping context, SICI is greater than go trials after the stop signal (Coxon et al., 2006; Hermans et al., 2019) and negatively associated with stop-signal reaction time, even when measured at rest (Chowdhury et al., 2019a, 2019b). In the current study, which implemented a selective stopping context, SICI in the left hand was decreased relative to signal onset during go trials but was maintained during partial-ignore_left_ and partial-stop_left_ trials. The maintenance of SICI during both partial-ignore and partial-stop trials suggests that the pause process likely drives SICI early after the partial signal. Interestingly, a release of SICI in the left hand was observed for partial-ignore_right_ but not partial-stop_right_ trials. It is possible that the pause-driven SICI effect was stronger or prolonged during partial-stop trials when stopping was explicitly required.

### 6.4 A pause-then-cancel model of selective stopping

The findings of the present study demonstrate the relevance of a pause-then-cancel model for selective stopping. Relying on the pause process could be advantageous during unimanual stopping since nonselective response inhibition would not impair the performance of other effectors (Wessel & Aron, 2017). In contrast, engaging a selective cancel process would provide for optimal selective stopping to not interfere with non-signalled effectors (Bissett & Logan, 2014). However, there was no evidence of a selective cancel process in the current study. Instead, response inhibition signatures were pause-driven during the critical period of response inhibition (150 to 250 ms) after the stop signal (Jana et al., 2020). It is possible that a selective cancel process could manifest later (e.g., Tatz et al., 2021), but time points beyond 250 ms reflect a period at which the response has already been executed in the non-signalled hand (MacDonald et al., 2014).

The amount of interference was twice as large during partial-stop than partial-ignore trials, despite the predominance of the pause process. Additional interference could result from a nonselective cancel process operating in parallel to the pause process (Diesburg & Wessel, 2021; Wadsley et al., 2022c). Alternatively, successful stopping may reflect instances where the pause process was stronger or prolonged compared to ignoring. Indeed, there was a null change from the onset in CME and SICI during partial-stop but inconclusive evidence for partial-ignore trials. In summary, the above findings indicate that response inhibition can be characterised by a nonselective pause from attentional capture, even in contexts that require selective stopping.

### 6.5 Limitations

The present study has some limitations. In-task TMS was delivered only at onset or 175 ms post-signal to collect a reliable number of stimulated trials per condition (Goldsworthy et al., 2016). The post-signal stimulation time was based on the expected period of the nonselective dip in CME during selective stopping (MacDonald et al., 2014). However, differences between partial-ignore and partial-stop trials may be discerned by evaluating CME modulation with a greater temporal resolution (e.g., Tatz et al., 2021).

Another limitation was the separation of EEG and TMS components, which were split so that the timing of TMS could be made participant-specific (e.g., Hermans et al., 2019) and to avoid excessive stimulation intensities due to coil displacement by EEG electrodes. However, combining EEG and TMS may better elucidate the functional relationship between β-bursts and SICI during selective stopping.

### 6.6 Conclusion

The present study aimed to determine how pause and cancel processes contribute to response inhibition during selective stopping. Response times in the non-signalled hand were delayed during partial trials, but overall interference was twice as large during partial-stop than partial-ignore trials. Stimulus-nonselective modulation occurred for β-bursts, CME, and SICI, irrespective of whether the assessed sensorimotor cortex was contralateral to the non-signalled hand. However, the magnitude of stopping-interference was not influenced by β-bursting. Overall, the findings indicate that nonselective response inhibition during selective stopping results from a nonselective pause process that cannot account for stopping-interference entirely.

## Acknowledgements

The authors thank April Ren and Federica Li Bassi for their assistance with data collection.

## References

Aron, A. R. (2011). From reactive to proactive and selective control: developing a richer model for stopping inappropriate responses. Biol Psychiatry, 69(12), e55–68. https://doi.org/10.1016/j.biopsych.2010.07.024

Aron, A. R., & Verbruggen, F. (2008). Stop the presses: dissociating a selective from a global mechanism for stopping. Psychol Sci, 19(11), 1146–1153. https://doi.org/10.1111/j.1467-9280.2008.02216.x

Awiszus, F., Borckardt, J.J. (2011). TMS Motor Threshold Assessment Tool. In (Version 2.0)

Bissett, P. G., & Logan, G. D. (2014). Selective stopping? Maybe not. J Exp Psychol Gen, 143(1), 455–472. https://doi.org/10.1037/a0032122

Chowdhury, N. S., Livesey, E. J., & Harris, J. A. (2019a). Contralateral and Ipsilateral Relationships between Intracortical Inhibition and Stopping Efficiency. Neuroscience, 415, 10–17. https://doi.org/10.1016/j.neuroscience.2019.07.013

Chowdhury, N. S., Livesey, E. J., & Harris, J. A. (2019b). Individual differences in intracortical inhibition during behavioural inhibition. Neuropsychologia, 124, 55–65. https://doi.org/10.1016/j.neuropsychologia.2019.01.008

Cowie, M. J., MacDonald, H. J., Cirillo, J., & Byblow, W. D. (2016). Proactive modulation of long-interval intracortical inhibition during response inhibition. J Neurophysiol, 116(2), 859–867. https://doi.org/10.1152/jn.00144.2016

Coxon, J. P., Stinear, C. M., & Byblow, W. D. (2006). Intracortical inhibition during volitional inhibition of prepared action. J Neurophysiol, 95(6), 3371–3383. https://doi.org/10.1152/jn.01334.2005

Coxon, J. P., Stinear, C. M., & Byblow, W. D. (2007). Selective inhibition of movement. J Neurophysiol, 97(3), 2480–2489. https://doi.org/10.1152/jn.01284.2006

De Jong, R., Coles, M. G., & Logan, G. D. (1995). Strategies and mechanisms in nonselective and selective inhibitory motor control. J Exp Psychol Hum Percept Perform, 21(3), 498–511. https://doi.org/10.1037//0096-1523.21.3.498

Delorme, A., & Makeig, S. (2004). EEGLAB: an open source toolbox for analysis of single-trial EEG dynamics including independent component analysis. J Neurosci Methods, 134(1), 9–21. https://doi.org/10.1016/j.jneumeth.2003.10.009

Diesburg, D. A., & Wessel, J. R. (2021). The Pause-then-Cancel model of human action-stopping: Theoretical considerations and empirical evidence. Neurosci Biobehav Rev, 129, 17–34. https://doi.org/10.1016/j.neubiorev.2021.07.019

Duque, J., Greenhouse, I., Labruna, L., & Ivry, R. B. (2017). Physiological Markers of Motor Inhibition during Human Behavior. Trends Neurosci, 40(4), 219–236. https://doi.org/10.1016/j.tins.2017.02.006

Enz, N., Ruddy, K. L., Rueda-Delgado, L. M., & Whelan, R. (2021). Volume of beta-Bursts, But Not Their Rate, Predicts Successful Response Inhibition. J Neurosci, 41(23), 5069–5079. https://doi.org/10.1523/JNEUROSCI.2231-20.2021

Goldsworthy, M. R., Hordacre, B., & Ridding, M. C. (2016). Minimum number of trials required for within- and between-session reliability of TMS measures of corticospinal excitability. Neuroscience, 320, 205–209. https://doi.org/10.1016/j.neuroscience.2016.02.012

Hannah, R., Muralidharan, V., Sundby, K. K., & Aron, A. R. (2020). Temporally-precise disruption of prefrontal cortex informed by the timing of beta bursts impairs human action-stopping. Neuroimage, 222, 117222. https://doi.org/10.1016/j.neuroimage.2020.117222

Hermans, L., Maes, C., Pauwels, L., Cuypers, K., Heise, K. F., Swinnen, S. P., & Leunissen, I. (2019). Age-related alterations in the modulation of intracortical inhibition during stopping of actions. Aging (Albany NY), 11(2), 371–385. https://doi.org/10.18632/aging.101741

Iacullo, C., Diesburg, D. A., & Wessel, J. R. (2020). Non-selective inhibition of the motor system following unexpected and expected infrequent events. Exp Brain Res, 238(12), 2701–2710. https://doi.org/10.1007/s00221-020-05919-3

Jana, S., Hannah, R., Muralidharan, V., & Aron, A. R. (2020). Temporal cascade of frontal, motor and muscle processes underlying human action-stopping. Elife, 9. https://doi.org/10.7554/eLife.50371

JASP Team. (2020). JASP. In JASP Team. https://jasp-stats.org/

Kilavik, B. E., Zaepffel, M., Brovelli, A., MacKay, W. A., & Riehle, A. (2013). The ups and downs of beta oscillations in sensorimotor cortex. Exp Neurol, 245, 15–26. https://doi.org/10.1016/j.expneurol.2012.09.014

Ko, Y. T., & Miller, J. (2013). Signal-related contributions to stopping-interference effects in selective response inhibition. Exp Brain Res, 228(2), 205–212. https://doi.org/10.1007/s00221-013-3552-y

MacDonald, H. J., Coxon, J. P., Stinear, C. M., & Byblow, W. D. (2014). The fall and rise of corticomotor excitability with cancellation and reinitiation of prepared action. J Neurophysiol, 112(11), 2707–2717. https://doi.org/10.1152/jn.00366.2014

MacDonald, H. J., McMorland, A. J., Stinear, C. M., Coxon, J. P., & Byblow, W. D. (2017). An Activation Threshold Model for Response Inhibition. PLoS One, 12(1), e0169320. https://doi.org/10.1371/journal.pone.0169320

Muralidharan, V., Aron, A. R., & Schmidt, R. (2022). Transient beta modulates decision thresholds during human action-stopping. Neuroimage, 254, 119145. https://doi.org/10.1016/j.neuroimage.2022.119145

Peirce, J., Gray, J. R., Simpson, S., MacAskill, M., Hochenberger, R., Sogo, H., Kastman, E., & Lindelov, J. K. (2019). PsychoPy2: Experiments in behavior made easy. Behav Res Methods, 51(1), 195–203. https://doi.org/10.3758/s13428-018-01193-y

Peurala, S. H., Muller-Dahlhaus, J. F., Arai, N., & Ziemann, U. (2008). Interference of short-interval intracortical inhibition (SICI) and short-interval intracortical facilitation (SICF). Clin Neurophysiol, 119(10), 2291–2297. https://doi.org/10.1016/j.clinph.2008.05.031

Pion-Tonachini, L., Kreutz-Delgado, K., & Makeig, S. (2019). ICLabel: An automated electroencephalographic independent component classifier, dataset, and website. Neuroimage, 198, 181–197. https://doi.org/10.1016/j.neuroimage.2019.05.026

Raud, L., Huster, R. J., Ivry, R. B., Labruna, L., Messel, M. S., & Greenhouse, I. (2020). A Single Mechanism for Global and Selective Response Inhibition under the Influence of Motor Preparation. J Neurosci, 40(41), 7921–7935. https://doi.org/10.1523/JNEUROSCI.0607-20.2020

Sharp, D. J., Bonnelle, V., De Boissezon, X., Beckmann, C. F., James, S. G., Patel, M. C., & Mehta, M. A. (2010). Distinct frontal systems for response inhibition, attentional capture, and error processing. Proc Natl Acad Sci U S A, 107(13), 6106–6111. https://doi.org/10.1073/pnas.1000175107

Shin, H., Law, R., Tsutsui, S., Moore, C. I., & Jones, S. R. (2017). The rate of transient beta frequency events predicts behavior across tasks and species. Elife, 6. https://doi.org/10.7554/eLife.29086

Tatz, J. R., Soh, C., & Wessel, J. R. (2021). Common and Unique Inhibitory Control Signatures of Action-Stopping and Attentional Capture Suggest That Actions Are Stopped in Two Stages. J Neurosci, 41(42), 8826–8838. https://doi.org/10.1523/JNEUROSCI.1105-21.2021

Tenke, C. E., & Kayser, J. (2005). Reference-free quantification of EEG spectra: combining current source density (CSD) and frequency principal components analysis (fPCA). Clin Neurophysiol, 116(12), 2826–2846. https://doi.org/10.1016/j.clinph.2005.08.007

van den Bergh, D., Van Doorn, J., Marsman, M., Draws, T., Van Kesteren, E.-J., Derks, K., Dablander, F., Gronau, Q. F., Kucharský, Š., & Gupta, A. R. K. N. (2020). A tutorial on conducting and interpreting a Bayesian ANOVA in JASP. LAnnee psychologique, 120(1), 73–96.

van den Bergh, D., Wagenmakers, E.-J., & Aust, F. (2022). Bayesian Repeated-Measures ANOVA: An Updated Methodology Implemented in JASP. PsyArXiv. https://doi.org/10.31234/osf.io/fb8zn.

van Doorn, J., van den Bergh, D., Bohm, U., Dablander, F., Derks, K., Draws, T., Etz, A., Evans, N. J., Gronau, Q. F., Haaf, J. M., Hinne, M., Kucharsky, S., Ly, A., Marsman, M., Matzke, D., Gupta, A., Sarafoglou, A., Stefan, A., Voelkel, J. G., & Wagenmakers, E. J. (2021). The JASP guidelines for conducting and reporting a Bayesian analysis. Psychon Bull Rev, 28(3), 813–826. https://doi.org/10.3758/s13423-020-01798-5

Veale, J. F. (2014). Edinburgh Handedness Inventory - Short Form: a revised version based on confirmatory factor analysis. Laterality, 19(2), 164–177. https://doi.org/10.1080/1357650X.2013.783045

Verbruggen, F., Chambers, C. D., & Logan, G. D. (2013). Fictitious inhibitory differences: how skewness and slowing distort the estimation of stopping latencies. Psychol Sci, 24(3), 352–362. https://doi.org/10.1177/0956797612457390

Wadsley, C. G., Cirillo, J., Nieuwenhuys, A., & Byblow, W. D. (2022a). Comparing anticipatory and stop-signal response inhibition with a novel, open-source selective stopping toolbox. PsyArXiv.

Wadsley, C. G., Cirillo, J., Nieuwenhuys, A., & Byblow, W. D. (2022b). Decoupling countermands nonselective response inhibition during selective stopping. J Neurophysiol, 127(1), 188–203. https://doi.org/10.1152/jn.00495.2021

Wadsley, C. G., Cirillo, J., Nieuwenhuys, A., & Byblow, W. D. (2022c). Stopping Interference in Response Inhibition: Behavioral and Neural Signatures of Selective Stopping. J Neurosci, 42(2), 156–165. https://doi.org/10.1523/JNEUROSCI.0668-21.2021

Wessel, J. R. (2020). beta-Bursts Reveal the Trial-to-Trial Dynamics of Movement Initiation and Cancellation. J Neurosci, 40(2), 411–423. https://doi.org/10.1523/JNEUROSCI.1887-19.2019

Wessel, J. R., & Aron, A. R. (2013). Unexpected events induce motor slowing via a brain mechanism for action-stopping with global suppressive effects. J Neurosci, 33(47), 18481–18491. https://doi.org/10.1523/JNEUROSCI.3456-13.2013

Wessel, J. R., & Aron, A. R. (2017). On the Globality of Motor Suppression: Unexpected Events and Their Influence on Behavior and Cognition. Neuron, 93(2), 259–280. https://doi.org/10.1016/j.neuron.2016.12.013

Westfall, P. H. (1997). Multiple Testing of General Contrasts Using Logical Constraints and Correlations. Journal of the American Statistical Association, 92(437), 299–306. https://doi.org/10.1080/01621459.1997.10473627

Wilcox, R. R. (2010). Fundamentals of Modern Statistical Methods (2 ed.). Springer-Verlag New York. https://doi.org/10.1007/978-1-4419-5525-8

Xu, J., Westrick, Z., & Ivry, R. B. (2015). Selective inhibition of a multicomponent response can be achieved without cost. J Neurophysiol, 113(2), 455–465. https://doi.org/10.1152/jn.00101.2014

